# DNA methylation aging and transcriptomic studies in horses

**DOI:** 10.1101/2021.03.11.435032

**Authors:** Steve Horvath, Amin Haghani, Sichong Peng, Erin N. Hales, Joseph A. Zoller, Ken Raj, Brenda Larison, Jessica L. Petersen, Rebecca R. Bellone, Carrie J. Finno

## Abstract

Human DNA methylation profiles have been used successfully to develop highly accurate biomarkers of aging (“epigenetic clocks”). Here, we describe epigenetic clocks for horses, based on methylation profiles of CpGs with flanking DNA sequences that are highly conserved between multiple mammalian species. Methylation levels of these CpGs were measured using a custom-designed Infinium array (HorvathMammalMethylChip40). We generated 336 DNA methylation profiles from 42 different horse tissues and body parts, which we used to develop five epigenetic clocks for horses: a multi-tissue clock, a blood clock, a liver clock and two dual-species clocks that apply to both horses and humans. Epigenetic age measured by these clocks show that while castration affects the basal methylation levels of individual cytosines, it does not exert a significant impact on the epigenetic aging rate of the horse. We observed that most age-related CpGs are adjacent to developmental genes. Consistently, these CpGs reside in bivalent chromatin domains and polycomb repressive targets, which are elements that control expression of developmental genes. The availability of an RNA expression atlas of these tissues allowed us to correlate CpG methylation, their corresponding contextual chromatin features and gene expression. This analysis revealed that while increased methylation of CpGs in enhancers is likely to repress gene expression, methylation of CpGs in bivalent chromatin domains on the other hand is likely to stimulate expression of the corresponding downstream loci, which are often developmental genes. This supports the notion that aging may be accompanied by increased expression of developmental genes. It is expected that the epigenetic clocks will be useful for identifying and validating anti-aging interventions for horses.

## INTRODUCTION

While there is a rich literature on human epigenetic clocks for individual and multiple tissues ^1-4^, we are not aware of any existing epigenetic clocks identified for horses. The human epigenetic clocks have already found many biomedical applications, including the measure of biological age in human anti-aging clinical trials ^1,5^. This has instigated the development of similar clocks for other mammals such as mice and dogs as biomarkers for age related disease ^6-12^. Here, we aimed to develop and evaluate epigenetic clocks for horses and to specifically investigate changes in DNA methylation that accompany equine aging. The availability of corresponding RNA-seq data allowed for the correlation between e methylation changes and gene expression levels, which is an advancement over previous studies in other species.

It has long been known that the level of cellular DNA methylation changes with age ^13-15^. With the technical development of methylation arrays that profile large numbers of individual CpG positions in the genome, an opportunity arose to develop a highly accurate age-estimator for all human tissues ^1,2,4^. For example, the human pan-tissue clock combines the weighted average of methylation levels of 353 CpGs into an age estimate that is referred to as DNAm age or epigenetic age ^16^. While the human pan-tissue clock applies to chimpanzees, ^16^ it does not apply to more distantly related mammals as a result of evolutionary genome sequence divergence. Recently, others have constructed epigenetic clocks for mice ^6-11^. Overall, these publications indicate that the underlying biological principle of epigenetic clocks is shared between members of different mammalian species. In humans, the discrepancy between DNA methylation age and chronological age (which is termed “epigenetic age acceleration”) is predictive of multiple health conditions, even after adjusting for a variety of known risk factors ^17-22^. Specifically, epigenetic age acceleration is associated with, but not limited to, cognitive and physical functioning ^23^, Alzheimer’s disease ^24^, centenarian status ^21,25^, Down syndrome ^26^, progeria ^27,28^, HIV infection ^29^, Huntington’s disease ^30^, obesity ^31^, cancer ^32^, and menopause ^33^. Epigenetic age is also predictive of mortality, even after adjusting for known risk factors such as chronological age, sex, smoking status, and other risk factors ^17-22^. Collectively, the evidence is compelling that epigenetic age is an indicator of biological age ^34-36^.

To enhance our understanding of the mechanistic causes and consequences of epigenetic aging effects, it is useful to correlate age-related DNA methylation changes to gene expression in different tissues. To address this challenge, we generated DNA methylation profiles from across 42 horse tissues for which transcription profiles were available. We report here, the development of a DNA-methylation-based estimator of chronological age across the entire lifespan of horses and humans (dual species clocks). Next, we characterize age-related changes in methylation levels in horses. Finally, we correlate age-related cytosines to gene expression levels from neighboring genes across different horse tissues.

## Results

We generated DNA methylation profiles from various tissue samples from horses **(Table 1)**. Tissues other than blood and liver were collected from two mares used in the equine Functional Annotation of Animal Genomes (FAANG) initiative ^37^. Unsupervised hierarchical clustering analysis of these profiles led to distinct tissue-based clusters (color band in **Supplementary Figure 1**). A subsequent random forest analysis of sex and tissue type led to an error rate of 0 and 10%, according to the out-of-bag (OOB) estimates.

**Table 1.** Description of blood methylation data. N=Total number of samples per species. Number of females. Age: mean, minimum and maximum.

| Tissue | N | No. Female | Mean age | Min. Age | Max. Age |
| --- | --- | --- | --- | --- | --- |
| Adipose | 2 | 2 | 4.5 | 4 | 5 |
| AdrenalCortex | 2 | 2 | 4.5 | 4 | 5 |
| Blood | 188 | 129 | 11.2 | 0.005 | 28 |
| Cartilage | 2 | 2 | 4.5 | 4 | 5 |
| Cecum | 4 | 4 | 4.5 | 4 | 5 |
| Cerebellum | 4 | 4 | 4.5 | 4 | 5 |
| Cortex | 2 | 2 | 4.5 | 4 | 5 |
| Duodenum | 2 | 2 | 4.5 | 4 | 5 |
| Fibroblast | 2 | 2 | 4.5 | 4 | 5 |
| Heart | 8 | 8 | 4.5 | 4 | 5 |
| Hypothalamus | 4 | 4 | 4.5 | 4 | 5 |
| Ileum | 2 | 2 | 4.5 | 4 | 5 |
| Jejunum | 2 | 2 | 4.5 | 4 | 5 |
| Keratinocyte | 2 | 2 | 4.5 | 4 | 5 |
| Kidney | 6 | 6 | 4.5 | 4 | 5 |
| Lamina | 2 | 2 | 4.5 | 4 | 5 |
| Larynx | 2 | 2 | 4.5 | 4 | 5 |
| Liver | 48 | 24 | 4.42 | 0.042 | 29 |
| Lung | 2 | 2 | 4.5 | 4 | 5 |
| Mammary | 4 | 4 | 4.5 | 4 | 5 |
| MitralValve | 2 | 2 | 4.5 | 4 | 5 |
| Muscle | 6 | 6 | 4.5 | 4 | 5 |
| OccipitalCortex | 2 | 2 | 4.5 | 4 | 5 |
| Ovary | 2 | 2 | 4.5 | 4 | 5 |
| ParietalCortex | 4 | 4 | 4.5 | 4 | 5 |
| Pituitary | 4 | 4 | 4.5 | 4 | 5 |
| Sacrocaudalis | 2 | 2 | 4.5 | 4 | 5 |
| Skin | 2 | 2 | 4.5 | 4 | 5 |
| SpinalCord | 4 | 4 | 4.5 | 4 | 5 |
| Spleen | 2 | 2 | 4.5 | 4 | 5 |
| Suspensory | 2 | 2 | 4.5 | 4 | 5 |
| TemporalCortex | 2 | 2 | 4.5 | 4 | 5 |
| Tendon | 4 | 4 | 4.5 | 4 | 5 |
| Uterus | 2 | 2 | 4.5 | 4 | 5 |

**Figure 1:**
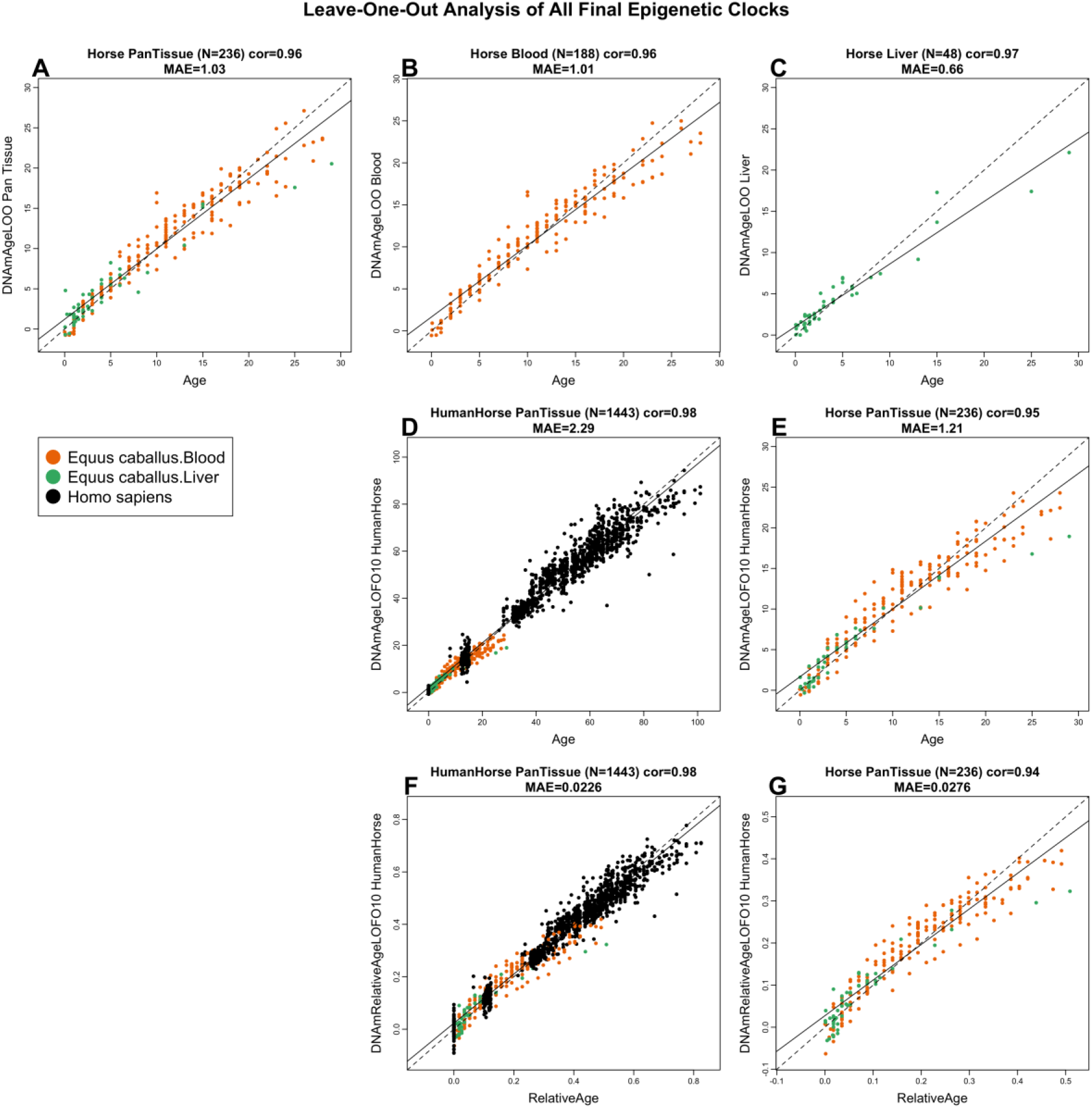
Cross-validation study of epigenetic clocks for horses and humans. Chronological age (x-axis) versus leave-one-sample-out (LOSO) estimate of DNA methylation age (y-axis, in units of years) for A) the multi-tissue clock for horse blood and liver, B) horse blood clock, C) horse liver clock. D) Ten-fold cross validation (LOFO10) analysis of the human-horse clock for chronological age. Dots are colored by species (black=human) and horse tissue type (green=liver, orange=blood). E) Same as panel D) but restricted to horses. F) Ten-fold cross validation analysis of the human-horse clock for relative age, which is the ratio of chronological age to the maximum recorded lifespan of the respective species. G) Same as panel D) but restricted to horses. Each panel reports sample size, correlation coefficient, median absolute error (MAE).

### Epigenetic clocks

We developed five epigenetic clocks for horses that vary with regards to three properties: species, tissue type, and measure of age. The multi-tissue horse clock was trained on horse blood and liver DNA methylation profiles, while the dual-species (human-horse) epigenetic clocks were trained using horse and human DNA methylation data. The resulting two human-horse clocks mutually differ by way of age measurement. One estimates *chronological age*s of horses and humans (in units of years), while the other estimates *relative* age, which is the ratio of chronological age of an animal to the maximum lifespan of its species (122.5 for humans and 57 for horses, Methods). The relative age ratio (with resulting values between 0 and 1) allows alignment and biologically meaningful comparison between species with different lifespans, which cannot otherwise be afforded by direct comparison of their chronological ages.

To arrive at unbiased estimates of the epigenetic clocks, we carried out cross-validation analysis of the training data. To develop the horse clocks, the training data employed consisted of horse blood and/or liver DNA methylation profiles, while human and horse DNA methylation profiles constituted the training data for both the human-horse clocks. The cross-validation study reports unbiased estimates of the age correlation R (defined as Pearson correlation between chronological age and its estimate, DNAm age), as well as the median absolute error. As indicated by its name, the horse multi-tissue clock is highly accurate in age estimation of blood and liver (R=0.96 and median absolute error 1.0 years, **Figure 1A**). The horse clocks for blood and liver samples lead to similar level of high accuracy (**Figure 1B,C**).

The human-horse clock for chronological age is highly accurate when DNA methylation profiles of both species are analyzed together (R=0.98, **Figure 1D**), and remains remarkably accurate when restricted to horse blood and liver samples (R=0.95, **Figure 1E**). Similarly, the human-horse clock for *relative age* exhibits high correlation regardless of whether the analysis is applied to samples from both species (R=0.98, **Figure 1F**) or only to horse samples (R=0.94, **Figure 1G**). This demonstrates that relative age circumvents the skewing that is inherent when chronological age of species with different lifespans is measured using a single formula.

### Horse clocks applied to plains zebras

Next, we applied the five horse clocks to blood and biopsy (skin) samples from plains zebras. According to the five different horse clocks, the DNAmAge estimates from zebra blood correlate highly with the age of the zebra (**Figure 2**). The results are less impressive for skin samples from zebras where horse clocks either over-or underestimate the chronological ages of zebras. Overall, these results show that the horse clocks can be applied to blood samples but not to skin samples from zebras.

**Figure 2.**
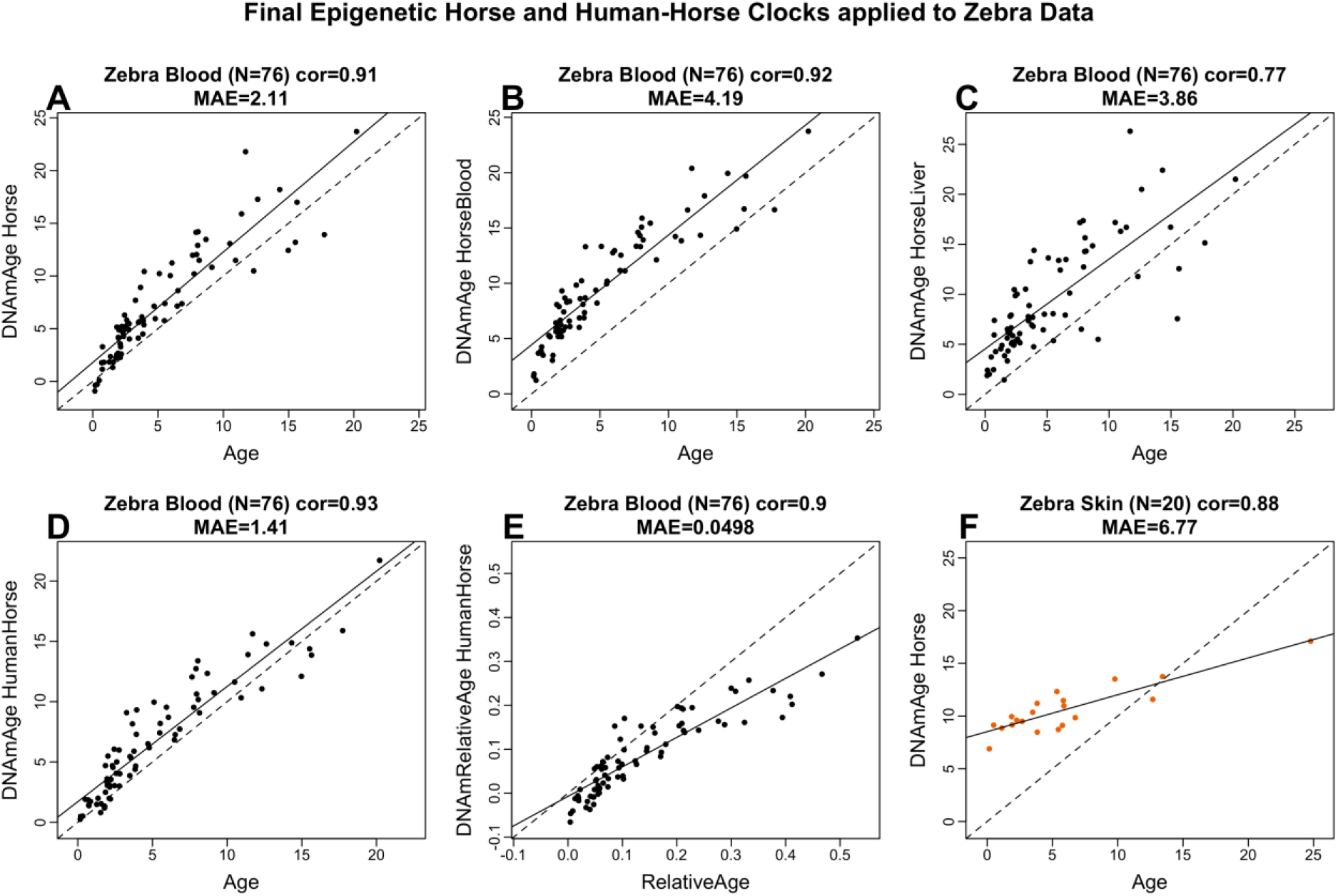
Horse clocks applied to blood and skin samples from plains zebra. Each panel corresponds to a different horse clock: A) Multi-tissue clock, B) blood clock, C) liver clock, D) human-horse clock for chronological age, E) human-horse clock for relative age F) Multi-tissue horse clock applied to skin samples from zebras. The solid line corresponds to linear least squares regression and the diagonal line to y=x.

### EWAS of age in horse tissues

The mammalian methylation array contains 31,836 probes that could be aligned to specific loci adjacent to 5093 unique genes in horse (Equus caballus.EquCab3.0.100) genome. Our epigenome-wide association studies (EWAS) correlated each of these CpGs with chronological age in horse blood (n=188) and liver (n=48) samples. In total, at a nominal p-value < 10^−5^, 12,705 and 1,813 probes were found to be related to age in blood and liver, respectively. The top DNA methylation changes in each tissue are as follows: blood, *HOXC4* intron (z=20), and *NFIA* intron (z= -19); and liver, *IKZF4* exon (z=11), and upstream of *HMX3* (z= -10) (**Figure 3A; Table S1**). Tissue level meta-analysis identified 10,501 CpGs with strong age-related methylation changes in both blood and liver. Some of these include hypermethylation of cytosines close to *TMEM121B, LHFPL4* and *FOXD3* exons and *TBX18* promoter regions. In general, the genes proximal to age-related CpGs in horse are involved in developmental processes and includes polycomb repressor complex 2 (e.g. EED, SUZ12) targets that are marked with H3K27ME3 modification (**Figure S3; Table S5**). This is a general and conserved DNAm aging pattern in all mammalian species ^38^.

**Figure 3.**
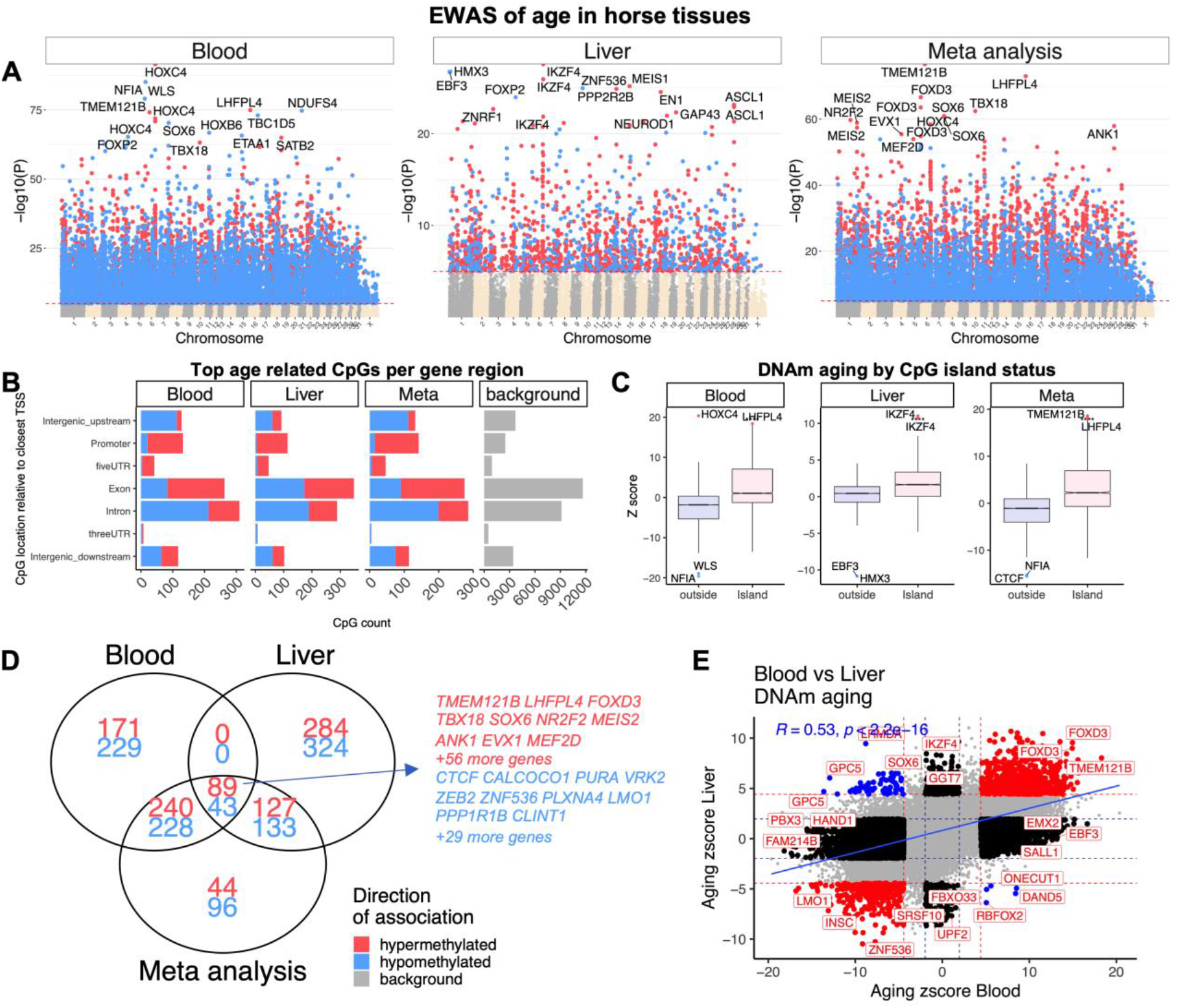
Epigenome-wide association (EWAS) of chronological age in horse. EWAS of age in blood (n=188), and liver (n=48) of horse. The meta-analysis results were calculated by Stouffer method. A) Manhattan plots of the EWAS of chronological age. The coordinates are estimated based on the alignment of Mammalian array probes to Equus_caballus.EquCab3.0.100 genome assembly. The direction of associations with p < 10^−5^(red dotted line) is highlighted by red (hypermethylated) and blue (hypomethylated) colors. Top 15 CpGs was labeled by the neighboring genes. B) Location of top CpGs in each tissue relative to adjacent transcriptional start sites. The grey color in the last panel represents the genomic location of 31836 mammalian methylation array probes mapped to the horse genome. C) Box plot analysis of position of age-associated CpGs in reference to CpG islands. The top age-related CpGs in each tissue are labeled by adjacent genes. **** p<10^−4^. D) Venn diagram of the overlap of the top 1000 (500 in each direction) significant CpGs in each analysis. E) Sector plot of DNA methylation aging in blood and liver of horse tissues. Red dotted line: p<10^−4^; blue dotted line: p>0.05; Red dots: shared CpGs; black dots: tissue specific changes; blue dots: CpGs whose age correlation differs between blood and liver tissue.

Age-related CpGs were found to be located in all genic and intergenic regions that can be defined relative to transcriptional start sites (**Figure 3B**). As expected, CpGs in regulatory regions, including promoters and 5’UTR, were mainly hypermethylated with age. This result paralleled a higher positive association of CpG islands with age than non-island CpGs in horse tissues (**Figure 3C**). The analysis also showed a partial overlap of DNAm aging between blood and liver (**Figure 3D**). These two horse tissues had a correlation of 0.5 for age-related CpG (**Figure 3E**). Although a large number of CpGs were changed by age in both liver and blood, there were several that were unique to each tissue. A subset of 26 CpGs were identified that had a divergent aging pattern between these two tissues. For example, while *GPC5* exon-1 is hypomethylated with age in blood, it is hypermethylated in horse liver. These divergent genes could be relevant in understanding developmental differences between blood and liver. Some of the shared and tissue specific DNAm changes are presented in **Figure S2**.

### Effect of castration

Castration is a common practice to modulate aggression of stallions. This presents an opportunity to test whether hormones, such as testosterone, the level of which is severely reduced in geldings, affects the rate of epigenetic aging. This has important implications to cancer risk in horses, as castrated males were previously shown to be at a higher risk for ocular SCC than females or stallions ^39-42^.We employed the horse clocks we developed to analyse the following subsets of the data: a) all male samples, b) males older than 5 years, and c) males younger than 15 years. We evaluated the effect of castration on the epigenetic age of blood and liver. Multivariate regression models that regressed leave-one-out estimates did not show a significant association between castration and aging, irrespective of the age stratum. Our multivariate analysis based on leave-one-sample out estimates has an obvious limitation: both castrated and intact animals were used in the training set which may condition out the effect of castration. Therefore, we repeated the analysis by developing a clock in female samples (training data) and applying it to male samples (test data). Again, we did not find a significant association of castration on epigenetic aging in blood.

Aging effects on CpG methylation in geldings correlated highly with those of stallions (r=0.78) (**Figure 4B, C**). There were nevertheless a few loci with methylation levels that changed with age only in geldings. For example, a CpG in the exon of *FOXP2* is hypomethylated with age in geldings, while a CpG in the 3’UTR of *ABCA1* is hypomethylated only in stallions (**Figure 4D, E; Table S3**). In general, there was a moderate difference between geldings and stallions with regards to the baseline mean methylation of CpGs in blood (independent of age). Some of the top methylation signatures of geldings included hypermethylation in *RABAC1* intron, *PRPH* exon, and hypomethylation in *TRPS1* intron, and *AKAP6* intron (**Figure 4A; Table S2**). While this study indicates a modest castration effect on baseline DNA methylation level and age-related CpG methylation changes, the validation of this relationship requires a much larger sample size from all age ranges and also consideration for potential confounding effect of horse breeds. Castration-related genes, both with baseline or age differences, were related to development of nervous system, cartilage, connective tissue, and muscle physiology (**Figure S4; Table S6**).

**Figure 4.**
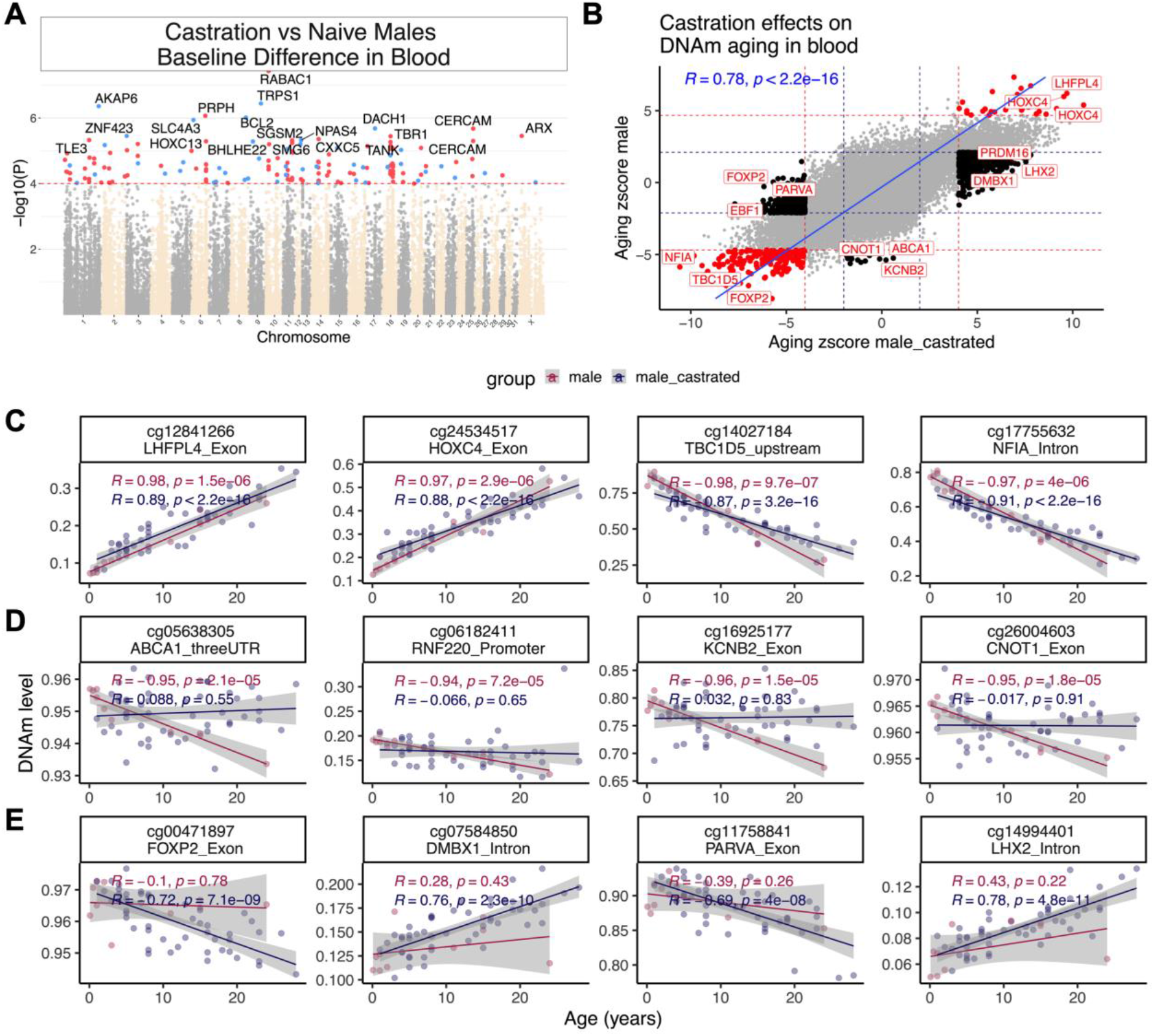
Castration moderately alters DNAm profile of horse blood. **A)** Manhattan plots of the EWAS of castration, in the blood of male horses. Co-variate: chronological age. Sample size: geldings, 48; stallions, 10. The coordinates are estimated based on the alignment of Mammalian array probes to Equus_caballus.EquCab3.0.100 genome assembly. The direction of associations with p < 10^−4^ (red dotted line) is highlighted by red (hypermethylated) and blue (hypomethylated) colors. Top 15 CpGs were labeled by the neighboring genes. B) Sector plot of DNA methylation aging by castration in blood of male horses. Red-dotted line: p<10^−4^; blue-dotted line: p>0.05; Red dots: age-related CpGs not affected by castration; black dots: CpGs whose aging pattern differs between geldings and stallions. Scatter plots of selected CpGs that change with age in both (C), or only stallions (D), or geldings (E) blood. The red dots and blue dots in the scatter plot correspond to blood samples from geldings and stallions, respectively.

### DNAm relate to gene expression differences in horse tissues

Meaningful interpretation of epigenetic findings requires the coupling of DNA methylation changes with those of gene expression. This challenge is further compounded by comparisons between tissues and species. Our study provides a rare opportunity to address this question by studying CpGs that are located in genomic regions that are conserved across mammalian species. Here, we integrated DNAm and RNAseq data from 57 samples (originating from 29 different tissues of two horses ^37^) to uncover the relationship between methylation changes of promoter CpGs with the expression of adjacent genes. Our analysis revealed that this relationship is dependent on the distance between the methylation site to the transcriptional start site (TSS) (**Figure 5A**). In general, methylation of CpGs that are closer to TSSs (from 10,000 nucleotides downstream to 1000 upstream of TSS) have a stronger repressive effect on mRNA levels (r= -0.2, p<2×10^−16^). This negative relationship was independent of CpG island status of the loci (**Figure S5**).

**Figure 5.**
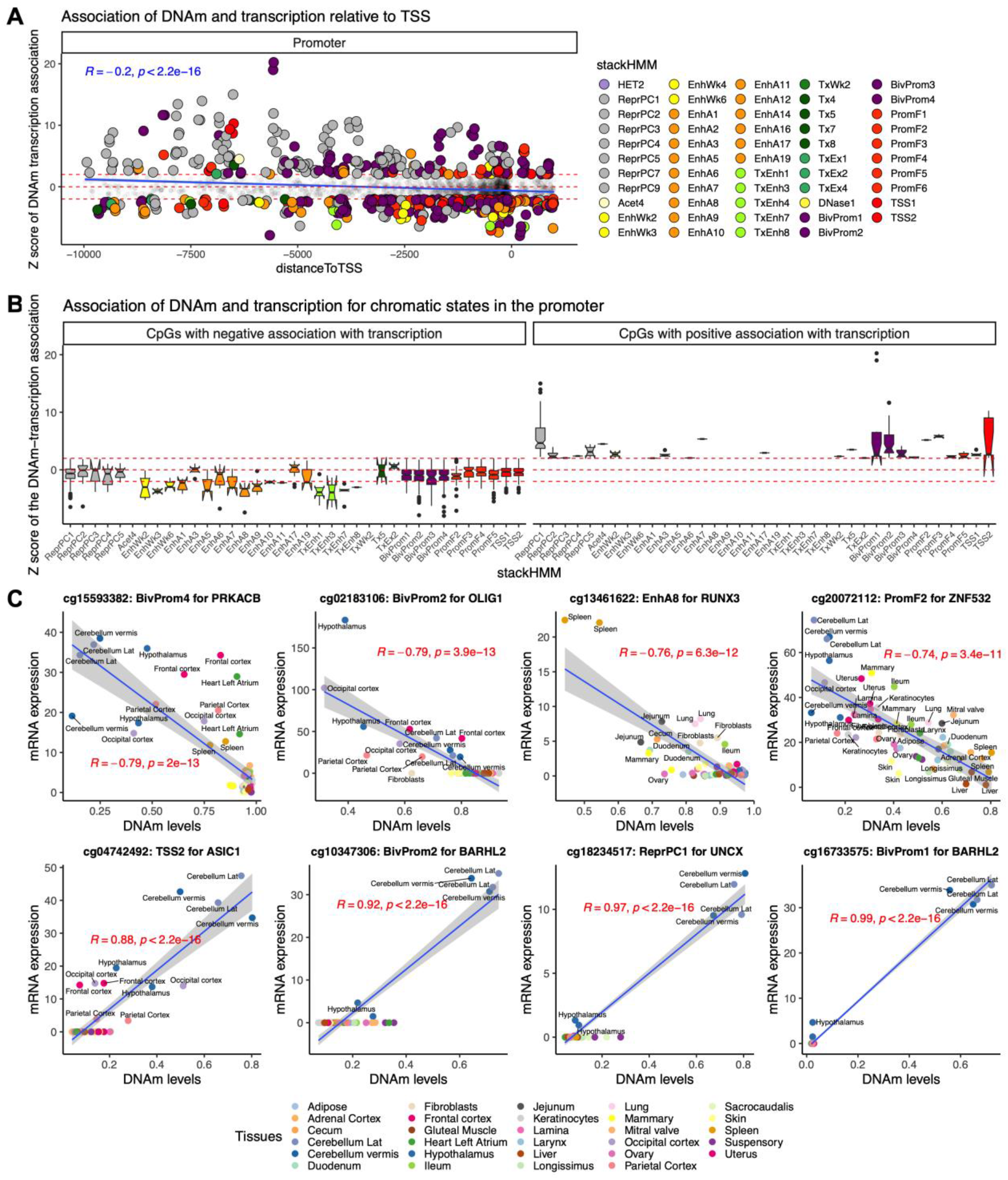
DNAm levels in promoters relate to gene expression changes in horse tissues. This analysis was based on linear correlation of DNAm and mRNA level of the adjacent genes in 29 tissues from the two female horses. A) Relationship of DNAm, mRNA expression, distance to transcription start site, and chromatin states in the gene promoters. The chromatin states are based on the stackHMM annotations, which represent a consensus chromatin state in over 100 human tissues ^43^. Red horizontal lines: z> 2 and z< -2 values. B) Boxplot of DNAm-mRNA association by stackHMM state in CpGs with significant cis-expression relationship. Only the stackHMM states with significant association (p<0.05) in any direction are shown in the figures. C) Scatter plots of selected CpGs with DNAm-mRNA association in horse tissues. Additional analysis by CpG island status and some sensitivity analyses are represented in the supplement (**Figure S5-S6**).

Regulation of gene expression however, is a multi-faceted process, of which methylation of the promoter is just one of the determinants. The chromatin context within which the CpGs are located is another feature that may exert a strong influence. As such, we first sought to ascertain the chromatin features within which the above CpGs are positioned, and then incorporate this information into the analysis of the impact of methylation of these CpGs on gene expression. Since chromatin states that are specific to the horse genome are presently unavailable, we used the “stacked chromatin states (stackHMM)” that identified chromatin features based on the consensus of over 100 human cell types ^43^. Despite the species difference, this approach can nevertheless prove highly informative because the design of the mammalian methylation array was based on DNA loci that are conserved across mammals ^44^, allowing chromatin features identified by stackHMM to be applied not only to the horse but other mammalian species as well. Interestingly, this analysis led to the observation that the contextual chromatin feature of CpGs was an even better indicator of the DNA methylation-gene expression relationship. In general, methylation of CpGs within enhancers appear to correlate with reduced gene expression (e.g. EnhWk2,3,6; EnhA3,6,7:9; and TxEnh1,3,7,8) (**Figure 5B, C)**. In contrast, increased gene expression is correlated with methylation of CpGs within polycomb repressed targets (e.g. ReprPC1,2,3,4,5), bivalent promoters (e.g. BivProm1-4), promoter flanks (e.g. PromF4,5), and transcriptional start sites (e.g. TSS1,2) (**Figure 5B, C)**. Correlations of individual CpG methylation to mRNA expression are reported in **Table S4**. Importantly, this interesting observation can soon be directly validated, as a horse-specific chromatin state annotation will become available as part of the on-going FAANG initiative ^45^.

## Discussion

We have previously developed several human epigenetic clocks from DNA methylation profiles that were derived from various versions of human Illumina DNA methylation arrays. As these arrays are specific to the human genome, their utility could not be extended to other species. A critical step toward crossing the species barrier was the design, development and use of a mammalian DNA methylation array that profiles 36,000 CpGs with flanking DNA sequences that are highly conserved across numerous mammalian species^44^. The employment of this array led to the acquisition of the most comprehensive epigenetic dataset of horses thus far. Using these data, we constructed highly accurate DNA methylation-based age estimators for horses that are applicable to their entire life course (from birth to old age). The successful derivation of a horse epigenetic clock using CpGs that are embedded within evolutionarily conserved DNA sequences across the mammalian class further confirms the conservation of the biological mechanisms that underpin aging.

The potential the horse clock could contribute to equine health is illustrated by the fact that human epigenetic age acceleration, as measured by the human clock, is associated with a wide array of age-related conditions. To accurately translate age-related findings from humans to horses (and vice-versa) requires a suitable measure of age-equivalence. We fulfilled this need through a two-step process. First, we combined DNA methylation profiles of horses and humans to generate a dual-species clock (human-horse), which is as accurate in estimating horse age as it is for human age, in the chronological unit of years. Second, we expressed the ages of every horse and human in ratios of the maximum recorded ages of their respective species (species lifespan). The mathematical operation of generating a ratio eliminates chronological unit of time and produces a value that indicates the age of the organism in respect to the maximum age of its own species. This allows a meaningful and realistic cross-species comparison of biological age. Collectively, the ability to use a single mathematical formula to measure epigenetic age of different species and the replacement of the chronological unit of time with proportion of lifespan, are two significant innovations that will propel cross-species research.

Beyond utilizing these methylation data sets to develop epigenetic clocks, we also investigated the characteristics of age-related CpGs and noted that the methylation of the majority of them were similar in direction in blood and liver, and that the genes that are proximal to them are largely developmental genes. This echoes the age-related CpG/genes identified across different tissues in other mammalian species. The connection between development and aging, albeit not immediately intuitive, is difficult to ignore. The enrichment of age-related CpGs to promoters, especially those with bivalent chromatin domains, and PRC2 targets suggest that changes of their methylation levels are likely to affect expression of downstream genes. It would indeed be very informative if such methylation changes could be correlated to gene expression changes. This has remained one of the challenging and limiting features of understanding DNA methylation changes because data for gene expression is often unavailable. Fortuitously, such data for these tissues are available in the horse. Here, we integrated DNAm and transcriptional data in a large horse tissue atlas. Our analysis suggests that cytosine methylation alone has a modest correlation with gene expression outcome. However, incorporation of contextual chromatin elements (enhancers, promoters etc.) to the analysis, increased the magnitude of correlation between CpG methylation and gene expression. Specifically, methylation of CpGs within enhancers are more likely to correlate with reduced gene expression, while methylation of CpGs in polycomb repressed targets, bivalent promoter, promoter flanks, and transcriptional start sites results in largely increased gene expression. This may at first appear counter-intuitive, as methylation of promoters is often associated with repression of transcription. It is to be noted however, that this notion is largely true for promoters with adjacent CpG islands. In the specific case of PRC targets and bivalent chromatin domains, our analysis is consistent with the recent observation that hypermethylation of bivalent chromatin domain results in reduced presence of the repressive histone H3K27me3, causing the balance of histone ratio towards the transcription-promoting H3K4me3 ^46^. This is consistent with the finding that DNA methylated regions of the genome are largely low in or devoid of H3K27me3, possibly due to the unfavorable binding of PRC2, which is required for methylation of H3K27 histone. Interestingly, methylation of bivalent chromatin domains was reported to correlate with increased expression of developmental genes ^47^, which incidentally are the predominant genes proximal to age related CpGs.

In addition to these valuable insights, this association between CpG methylation level, genomic elements and gene expression is a valuable tool to interpret methylation array findings from all mammalian species. For example, our analyses of age-related methylation changes in horse tissues reveals that CpGs that became increasingly methylated with age are located largely within promoters and CpG islands. Two of the top CpGs that were hypermethylated with age in horses included *TBX18* and *FOXD3* promoters. However, while the *TBX18* promoter had a strong negative correlation with DNAm-mRNA (z= -2.8), the *FOXD3* promoter showed a positive DNAm-mRNA correlation in horse tissues (z=1.5). Thus, we can deduce that *TBX18* expression decreases with age, but *FOXD3* mRNA levels will increase with age. Cross-species expansion of this finding is only possible due to the design of the mammalian methylation array for highly conserved genomic regions ^44^. Thus, this novel multi-omics analysis of horse tissues is a tool to link DNAm with transcription in other studies based on the mammalian methylation array. Such a link is essential for functional interpretation of the findings and also for the experimental design of gene-perturbation studies.

## Materials and Methods

### Study samples

#### Horses

We generated DNA methylation data from n=42 different horse tissues collected at necropsy (**Table 1**). The tissue atlas was generated from two Thoroughbred mares as part of the FAANG initiative ^37^, with the following tissues profiled: adipose (gluteal), adrenal cortex, blood (PBMCs; only n=1 mare), cartilage (only n=1 mare), cecum, cerebellum (2 samples each from lateral hemisphere and vermis), frontal cortex, duodenum, fibroblast, heart (2 samples each from the right atrium, left atrium, right ventricle, left ventricle), hypothalamus, ileum, jejunum, keratinocyte, kidney (kidney cortex and medulla), lamina, larynx (i.e. cricoarytenoideus dorsalis muscle), liver, lung, mammary gland, mitral valve of the heart, skeletal muscle (gluteal muscle and longissimus muscle), occipital cortex, ovary, parietal cortex, pituitary, sacrocaudalis dorsalis muscle, skin, spinal cord (C1 and T8), spleen, suspensory ligament, temporal cortex, tendon (deep digital flexor tendon and superficial digital flexor tendon), uterus ^37^. These tissues were also used for RNAseq analyses. This collection protocol was approved by the UC Davis Institutional Animal Care and Use Committee (Protocol#19037)

Blood samples were collected via venipuncture into EDTA tubes from across 24 different horse breeds (buffy coat). Most of the samples were from the Thoroughbred (TB) (n=79) and American Quarter Horse breeds (QH, n=62). For the following breeds, we had between one and six blood samples: Andalusian, Appaloosa, Arabian, Dutch Warmblood, Hanoverian, Holsteiner, Irish Sport Horse, Lipizzaner, Lusitano, mixed breed, Oldenburg, Paint or Paint cross, Percheron, Shire, Standardbred, Warmblood and Welsh Pony. The n=49 liver samples originated from necropsy collections of horses across 19 different breeds, with most of the liver samples from QHs (n=20). All collection protocols were approved by the UC Davis Institutional Animal Care and Use Committee (Protocols #20751 and 21455, respectively).

#### Plains zebras

The data from plains zebras (*Equus quagga*) are described in a companion paper (Larison et al 2021). Briefly, both blood (n=96) and biopsy (skin) (n=24) samples were obtained from a captive population of zebras maintained in a semi-wild state by the Quagga Project ^48^ in the Western Cape of South Africa. The population was founded in 1989 from 19 individuals (9 from Etosha National Park in Namibia, 10 from the Kwazulu-Natal in South Africa). Skin samples were taken by remote biopsy dart (1 mm wide by 20-25 mm deep plug) and preserved in RNAlater (Qiagen). Blood samples were taken opportunistically during veterinarian visits and preserved in EDTA tubes. Most samples were collected from different individuals, except for two animals that were sampled twice some years apart. All samples were stored at -20 °C. Samples were collected under a protocol approved by the Research Safety and Animal Welfare Administration, University of California Los Angeles: ARC # 2009-090-31, originally approved in 2009. After eliminating samples with low confidence for individual identity and age, we retained 76 blood samples, and 20 skin samples. We retained the founder, however, in an effort to extend the age range represented in the skin clock.

#### Human tissue samples

To construct the human-horse clock, we analyzed the generated methylation data from n=1,207 human tissue samples (adipose, blood, bone marrow, dermis, epidermis, heart, keratinocytes, fibroblasts, kidney, liver, lung, lymph node, muscle, pituitary, skin, spleen) from individuals whose ages ranged from 0 to 93. The tissue samples came from three sources: tissue and organ samples from the National NeuroAIDS Tissue Consortium ^49^, blood samples from the Cape Town Adolescent Antiretroviral Cohort study ^50^, and blood, skin and other primary cells provided by Kenneth Raj ^51^. Ethics approval was provided for all studies (IRB#15-001454, IRB#16-000471, IRB#18-000315, IRB#16-002028).

### DNA methylation data

The mammalian DNA methylation arrays were profiled using a custom Infinium methylation array (HorvathMammalMethylChip40) based on 37492 CpG sites. Out of these sites, 1951 were selected based on their utility for human biomarker studies. These CpGs, which were previously implemented in human Illumina Infinium arrays (EPIC, 450K), were selected due to their relevance for estimating age, blood cell counts or the proportion of neurons in brain tissue. The remaining 35541 probes were chosen to assess cytosine DNA methylation levels in mammalian species ^44^. The particular subset of species for each probe is provided in the chip manifest file can be found at Gene Expression Omnibus (GEO) at NCBI as platform GPL28271. The SeSaMe normalization method was used to define beta values for each probe ^52^. This study focused on 31,836 CpGs that could be mapped to the horse genome.

### RNA seq data

Strand-specific RNA libraries were created following poly-A selection. Libraries were sequenced at 2×150bp on Illumina HiSeq2500, with a targeted depth of 30 million reads. RNAseq data were used to quantify transcripts annotated in Ensemble annotation (GCA_002863925.1, release 103) using Salmon mapping-based mode ^53^.

### Penalized Regression models

Penalized regression models were created with glmnet ^54^. We investigated models produced by “elastic net” regression (alpha=0.5). The optimal penalty parameters in all cases were determined automatically by using a 10 fold internal cross-validation (cv.glmnet) on the training set.

We performed a cross-validation scheme for arriving at unbiased estimates of the accuracy of the different DNAm based age estimators. One type consisted of leaving out a single sample (LOOCV) from the regression, predicting an age for that sample, and iterating over all samples. A critical step is the transformation of chronological age (the dependent variable). While no transformation was used for the blood clock for horses, we did use a log linear transformation for the dual species clock of chronological age (**Supplement**). Details on the clocks (CpGs, genome coordinates) and R software code are provided in the Supplement

### Relative age estimation

Relative age estimation was performed to introduce biological meaning into the age estimates of horses and humans, which have very different lifespans. Additionally, this estimation serves to overcome the inevitable skewing due to unequal distribution of data points from horses and humans across age range. Relative age estimations were calculated using the formula: Relative age= Age/maxLifespan where the maximum lifespan for the two species was chosen from the “anAge” database (57 for horses and 122.5 for humans ^55^). The maximum for horses (57 years) will sound too high for many experts but miniature horses appear to live longer. The anAge data base ^55,56^ record for horse states the following. Quote “*One Icelandic miniature horse named …is reported to have lived 57 years (Richard Miller, pers. comm*.*). Anecdotal evidence tells of a horse…that lived for 62 years in England, but that record is unverified*.” The oldest regular sized horse appears to have reached an age of 53 according to the Official Guide for Determining the Age of the Horse from American Association of Equine Practitioners (B. Wright 1999 URL).

### Epigenome wide association studies of age

EWAS was performed in each tissue separately using the R function “standardScreeningNumericTrait” from the “WGCNA” R package^57^. Next the results were combined across tissues using Stouffer’s meta-analysis method.

### DNAm-mRNA integration

The mammalian methylation array carefully characterized the location of all target CpGs relative to the adjacent transcriptional start site in horse genome ^44^. Thus, we could link each CpG with mRNA level of the adjacent gene in horse tissue atlas. The associations were analyzed by the R function “standardScreeningNumericTrait” from the “WGCNA” R package ^57^.

## URL

B. Wright (1999) Official Guide for Determining the Age of the Horse, American Association of Equine Practitioners. http://www.omafra.gov.on.ca/english/livestock/horses/facts/info_age.htm

## Data availability

The RNA-seq data can be downloaded from https://www.ebi.ac.uk/ena/data/view/ERA1487553

The methylation data will be made publicly available as part of the data release from the Mammalian Methylation Array Consortium. Genome annotations of these CpGs can be found on Github https://github.com/shorvath/MammalianMethylationConsortium

## Acknowledgements

This work was supported by the Paul G. Allen Frontiers Group (SH). Funding for horse tissue sample collection was provided by the Grayson Jockey Club Foundation, USDA NRSP-8 and the UC Davis Center for Equine Health. Support for C.J.F. was provided by the National Institutes of Health (NIH) (K01OD015134 and L40 TR001136).

