## Supplementary Figures and Tables for "DNA methylation aging and transcriptomic studies in horses"

### SUPPLEMENTARY MATERIAL

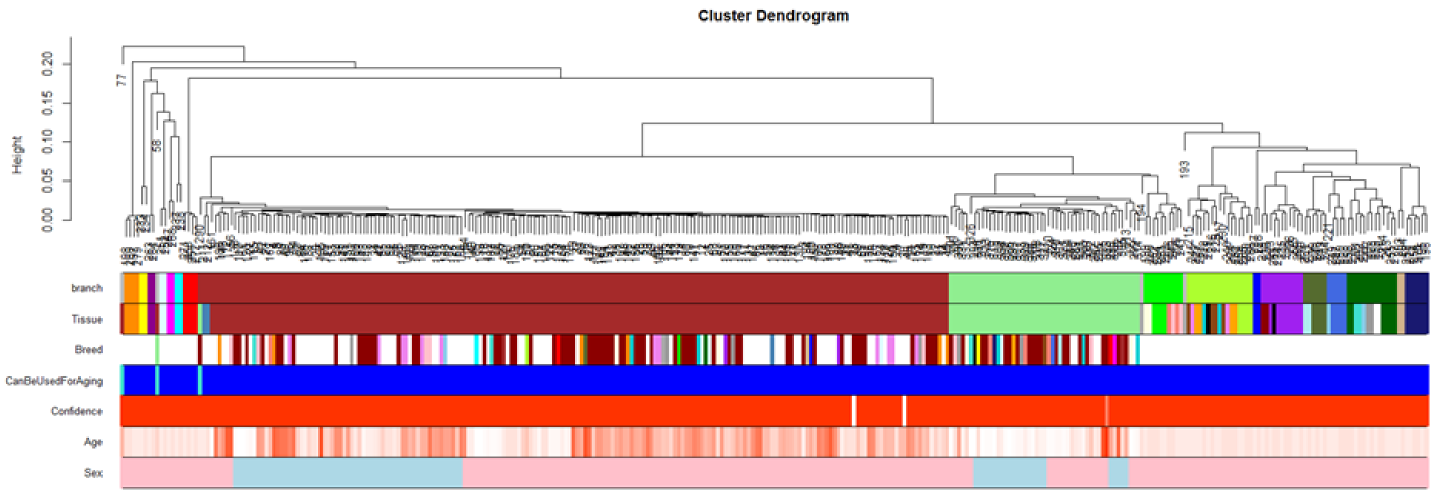


**Supplementary Figure 1. Unsupervised hierarchical clustering of blood samples from horses** Average linkage hierarchical clustering based on the inter-array correlation coefficient (Pearson correlation). The cluster branches (first color band) correspond to tissue type (second color band).

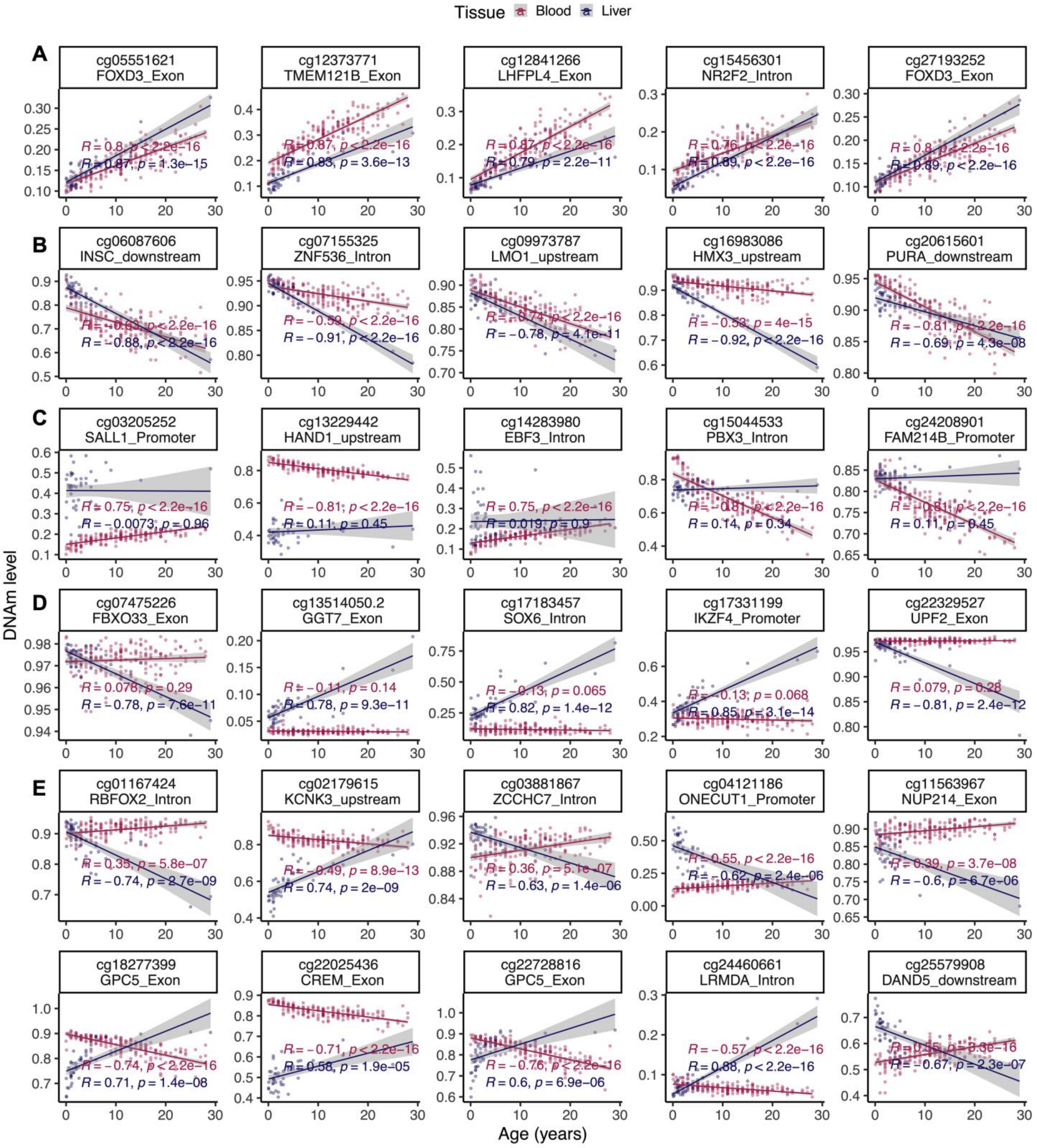


Figure S2. Scatter plots of age-related changes in selected CpGs in blood and liver of horse. A) CpGs that are hypermethyalted with age in both blood and liver. B) CpGs that are hypomethylated with age in both blood and liver. C) Examples of blood specific changes. D) Examples of liver specific changes. E) Selected CpGs with divergent aging pattern between blood and liver of horse.

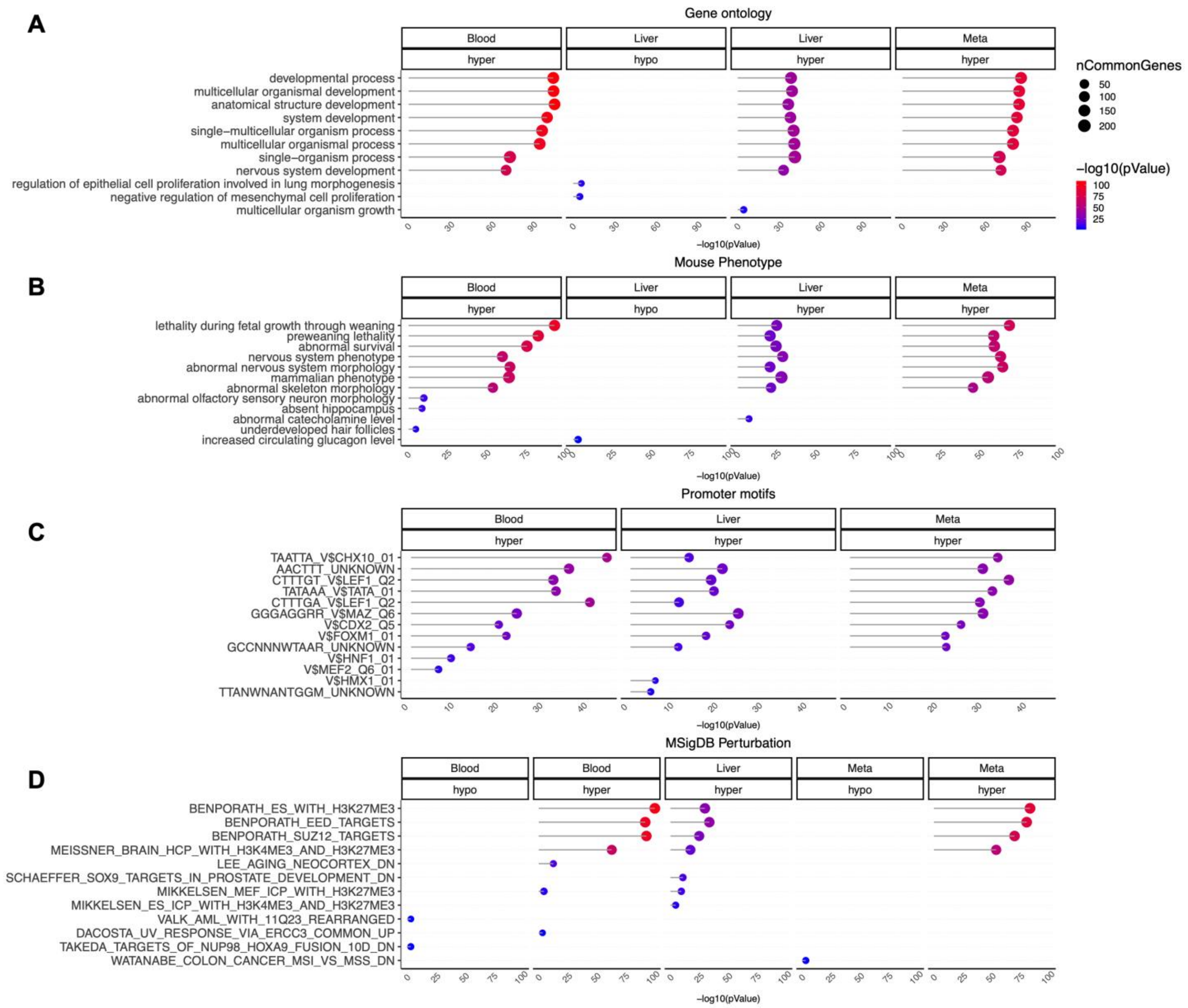


**Figure S3. Gene set enrichment analysis of DNA methylation aging in different horse tissues**. The gene level enrichment was done using GREAT analysis [1] and human Hg19 background. Datasets: gene ontology (A), mouse phenotypes (B), promoter motifs (C), and MSigDB Perturbation (D). The results were filtered for significance at $p < 10^{-5}$.

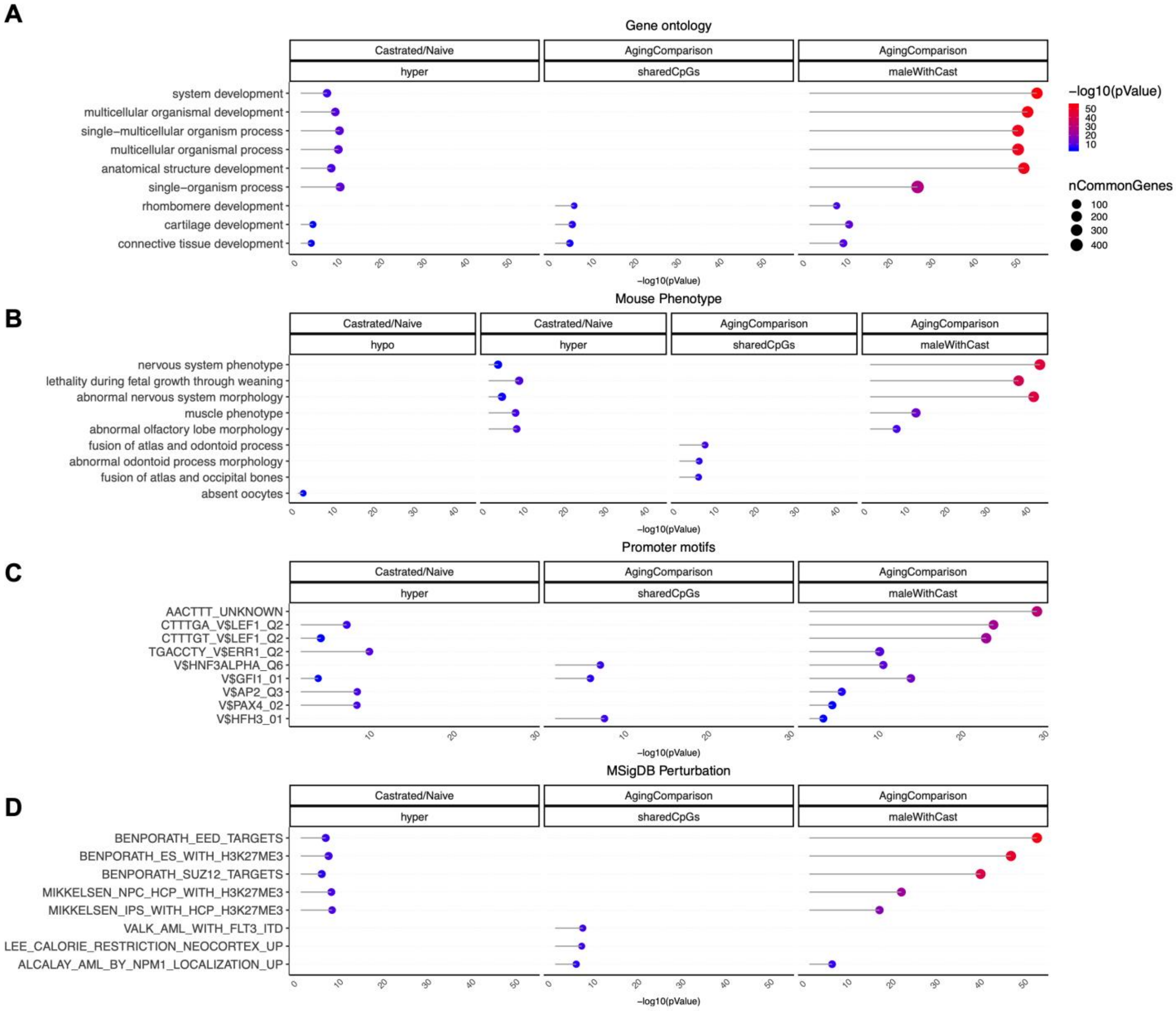


**Figure S4. Gene set enrichment analysis of DNA methylation changes by castration**. The gene level enrichment was done using GREAT analysis [1] and human Hg19 background. Datasets: gene ontology (A), mouse phenotypes (B), promoter motifs (C), and MSigDB Perturbation (D). The results were filtered for significance at $p < 10^{-3}$.

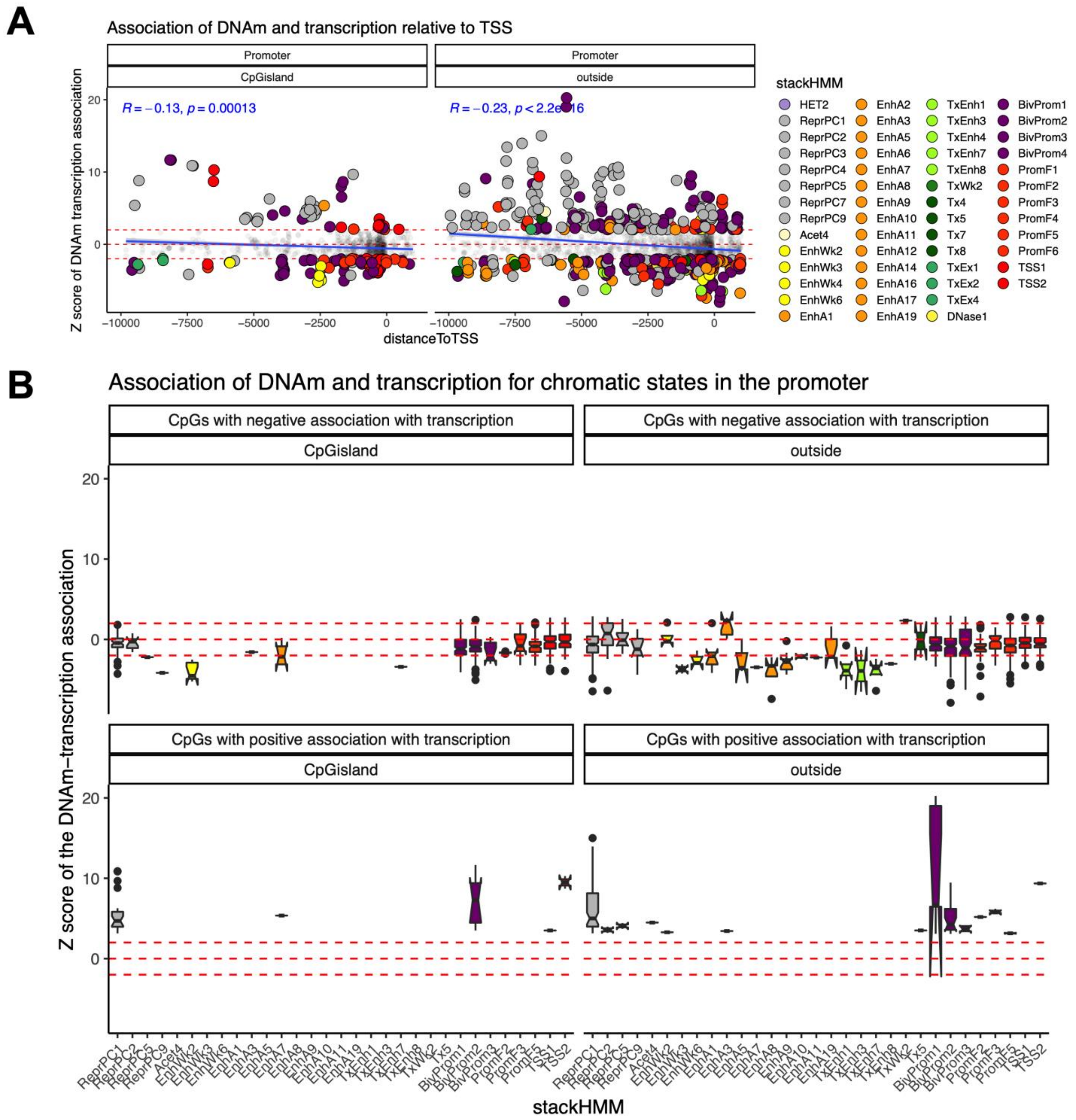


**Figure S5. Promoter CpG island status does not alter DNAm-mRNA associations.** A) Relationship of DNAm, mRNA expression, distance to transcription start site, and chromatin states in the gene promoters by CpG island status. The chromatin states are based on the stackHMM annotations, which represent a consensus chromatin state in over 100 human tissues [2]. Red horizontal lines: z> 2 and z< -2 values. B) Boxplot of DNAm-mRNA association by stackHMM state and CpG island status in CpGs with significant cis-expression relationship. Only the stackHMM states with significant association in any direction are shown in the figures. This analysis was based on linear correlation of DNAm and mRNA level of the adjacent genes in 29 tissues from the two horses.

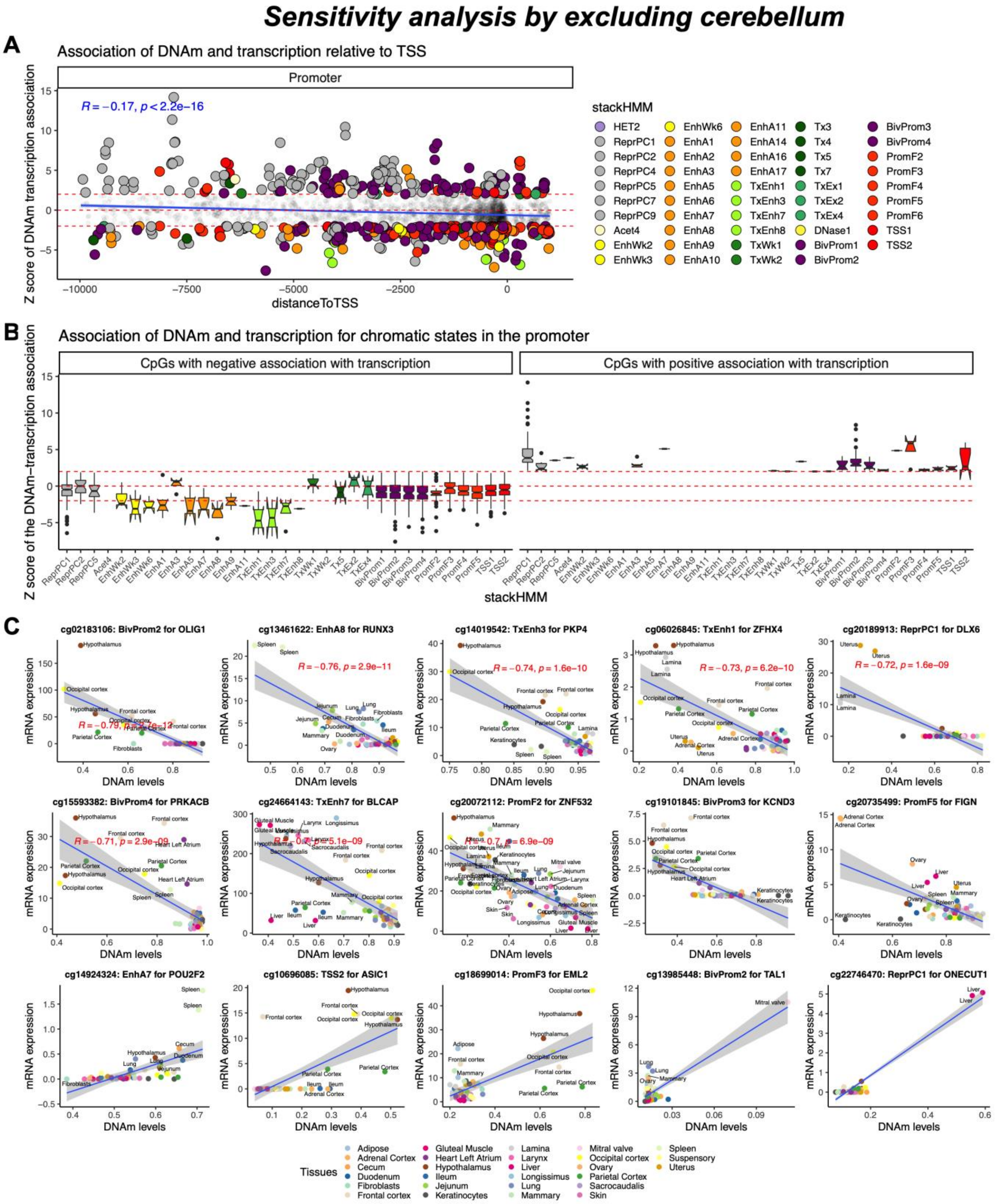


**Figure S6. Sensitivity of analysis DNAm-mRNA association.** Since cerebellum was a tissue with extreme DNAm-mRNA expression signatures in some stackHMM states, the cerebellum was excluded from the analysis. A) Negative association of distance to TSS with DNAm-mRNA expression association was not affected by excluding the cerebellum sample. B) Excluding the cerebellum did not affect stackHMM relationships with DNAm-mRNA changes. C) Scatter plots of selected CpGs with DNAm-mRNA association in horse tissues after excluding cerebellum from the analysis.

1 McLean, C. Y. *et al.* GREAT improves functional interpretation of cis-regulatory regions. *Nat Biotechnol* **28**, doi:10.1038/nbt.1630 (2010).
2 Vu, H. & Ernst, J. Universal annotation of the human genome through integration of over a thousand epigenomic datasets. *bioRxiv*, 2020.2011.2017.387134, doi:10.1101/2020.11.17.387134 (2020).
